# Diurnal plant and *Rhizophagus irregularis* transcriptional patterns are linked to shifts in cassava tissue partitioning in the field

**DOI:** 10.1101/2025.10.08.681086

**Authors:** Erica McGale, Chanz Robbins, Diego Camilo Peña-Quemba, Isabel Cristina Ceballos, Alia Rodriguez, Ian R. Sanders

## Abstract

- Cassava is a globally-important crop whose yields can radically increase with arbuscular mycorrhizal fungi (AMF) inoculation. However, extensive background noise in field environments makes it very challenging to understand how the cassava-AMF symbiosis confers benefits, which is especially important for future applications of AMF treatments.
- In two field experiments, we combined transcriptomics and allometric analyses to investigate functional variation in cassava-AMF interactions using sterile, single isolates of *Rhizophagus irregularis*. We developed a novel Index of Symbiotic Transcriptional Activity (ISTA) and accounted for sampling times to reduce transcriptomic noise and improve links to biomass traits.
- ISTA significantly correlated with cassava shoot biomass in an isolate-dependent manner, and allometric analyses revealed that *R. irregularis* isolates can either reinforce or uncouple cassava shoot-root relationships to maximize root yields. Differential expression and co-expression network analyses uncovered isolate-specific plant and fungal gene module responses. Including ISTA and sampling time as random effects enhanced detection of gene candidates, including down-regulated genes linked to higher yield.
- Our study uses novel and translatable transcriptomic tools to readily dissect variably field data, allowing new links to be found between AMF symbiotic functions and cassava yields.

## INTRODUCTION

Meeting the caloric needs of Earth’s nearly billion undernourished is of imminent importance (FAO, 2025). Issues with food availability are disproportionally abundant in sub-Saharan Africa, South America and South Asia (Azimi & Rahman, 2024; FAO, 2025), and will increase drastically with climate change (Sutton *et al*., 2013). In these same regions, *Manihot esculenta*, otherwise known as cassava, is a staple food crop with known importance for addressing food insecurity (Leff *et al*., 2004; Amelework *et al*., 2021). Cassava is the third most productive crop per unit area worldwide (FAO, 2018). However, its consumable products (i.e., tuberous roots) are belowground, making the success of its harvests difficult to predict. In addition, cultivars known to be high-performing in some locales have different growth and variation across field sites (Peña Venegas *et al*., 2021; Enesi *et al*., 2022). As of yet, aboveground traits of the cassava plant have not been shown to consistently predict belowground yields (Joaqui Barandica *et al*., 2016; Aliyu *et al*., 2019).

Applications of arbuscular mycorrhizal fungi (AMF) to agricultural soils can increase yields of crops including cassava (Ceballos *et al*., 2013; Zhang *et al*., 2019; Qin *et al*., 2022). Plants associating with AMF are thought to have higher access to soil nutrients and water through the fungus, as well as more stress tolerance (Bennett & Groten, 2022). The fungus receives carbon and fatty acids from the plant in return for conferred benefits (Shi *et al*., 2023). AMF from the Glomeraceae family are amongst the most abundant fungi found in tropical and sub-tropical soils (Ramírez Gómez, 2014; Séry *et al*., 2018; Thanni *et al*., 2022), emphasizing their availability and potential to address food insecurity in these areas. It has been shown that different isolates and single-spore progeny of the model Glomeraceae species, *Rhizophagus irregularis*, have considerable potential to alter cassava yields by up to 300% when applied in tropical and sub-tropical fields (Ceballos *et al*., 2013, 2019; Peña *et al*., 2020; Peña Venegas *et al*., 2021). However, more details on the mechanisms by which field AMF inoculations may increase crop yields are needed in order to make such applications more robust.

Previously, increased soil fertility (i.e., nutrient content) was thought to decrease the plant-AMF symbiosis, and thus, reduce the ability of inoculation to affect crop yield (Bender *et al*., 2019). Recent research has shown that increased soil nutrient content is not necessarily linked to root colonization percentage (Okon *et al*., 2010; Lutz *et al*., 2023), including in cassava (Peña Venegas *et al*., 2021). Additionally, percentages of colonized roots do not correlate with plant responsiveness to mycorrhization (Peña Venegas *et al*., 2021; Elliott *et al*., 2021; Lutz *et al*., 2023). Other environmental factors like pH and beneficial and pathogenic microbial biomass were shown to significantly contribute to the outcomes of AMF inoculation on crops (Herrmann *et al*., 2019; Lutz *et al*., 2023). Despite this emerging information, the plant and fungal pathways that are altered in response to AMF inoculation of crops, especially in plants with beneficial yield outcomes, are not well understood. A greater understanding of how changes in gene transcription in AMF inoculated cassava in the field may link to increases in cassava production is needed. Dissecting these plant-AMF effects in complex environments is essential for future application of AMF inoculums (French, 2017; Ferlian *et al*., 2018; Lekberg & Helgason, 2018). However, the immense difficulty of this task in the face of large amounts of variation in the field has prevented clear cassava-AMF field transcriptomic characterizations to date.

For cassava, differences in transcription linked to the plant’s development and tuberous root qualities were previously reported (Li *et al*., 2010; De Souza *et al*., 2017; Ding *et al*., 2025). These studies, however, did not consider the potential impact of AMF and its transcription, especially in realistic field situations. When applying an AMF inoculum in the field, it would be ideal to track the colonization and transcription of that specific isolate within plant roots. Then, the contribution of that isolate to changes in plant transcription, function, and yield, in the face of other factors, could be extrapolated. However, in practice, it is extremely difficult to assign transcription to a single AMF isolate under field conditions, where a community of different AMF exist in addition to the AMF added as inoculum. This would involve separating transcription by the inoculated AMF from that of the non-inoculated AMF already existing in field. Even with the ability to track colonization of a particular AMF species or genotype (Bodenhausen *et al*., 2021; Rodríguez-Yon *et al*., 2021; Heller & Carrara, 2022; Lee *et al*., 2024), limits still exist in determining the transcriptional activity of a single AMF. Whether inoculated AMF are producing new transcripts of particular genes, or indirectly influencing the entirety of the colonized AMF community to alter transcription of given genes is also extremely complex to disentangle (Sanders & Rodriguez, 2016). To begin advancing our understanding of cassava-AMF transcription with consequences on cassava yields in the field, we can analyze the total transcription of one model AMF species, like *R. irregularis*, in inoculated and non-inoculated field cassava roots. This can be matched to plant transcription, and innovative methods can be used to extract coordinated plant-AMF transcription from highly variable field data and explore its connection to previously unstudied effects on cassava yield outcomes.

We conducted two field experiments in Tauramena, Colombia, in consecutive years, with cassava inoculated with five genotypically distinct isolates of *R. irregularis* as well as six single-spore progeny derived from those isolates (Fig. 1A, B). We explored connections between cassava shoot and root biomass, total AMF colonization, and soil physical and chemical properties. We generated RNAseq data from the second field experiment and produced a plant transcriptional index to represent symbiotic activity. We tested if this index would better associate with cassava biomasses (Fig. 1C). We used two strategies to link plant and AMF gene transcription with increased cassava yield (root biomass) outcomes. First, we highlighted plant transcriptional modules that were altered especially when cassava plants, across inoculation treatments, outperformed non-inoculated plants. Second, we used allometric trajectories (Shingleton, 2010) to find shifts in cassava biomass partitioning linked to particular AMF isolates or single spore lines. We then determined plant and AMF transcription connected to distinguished shift types.

**Figure 1.**
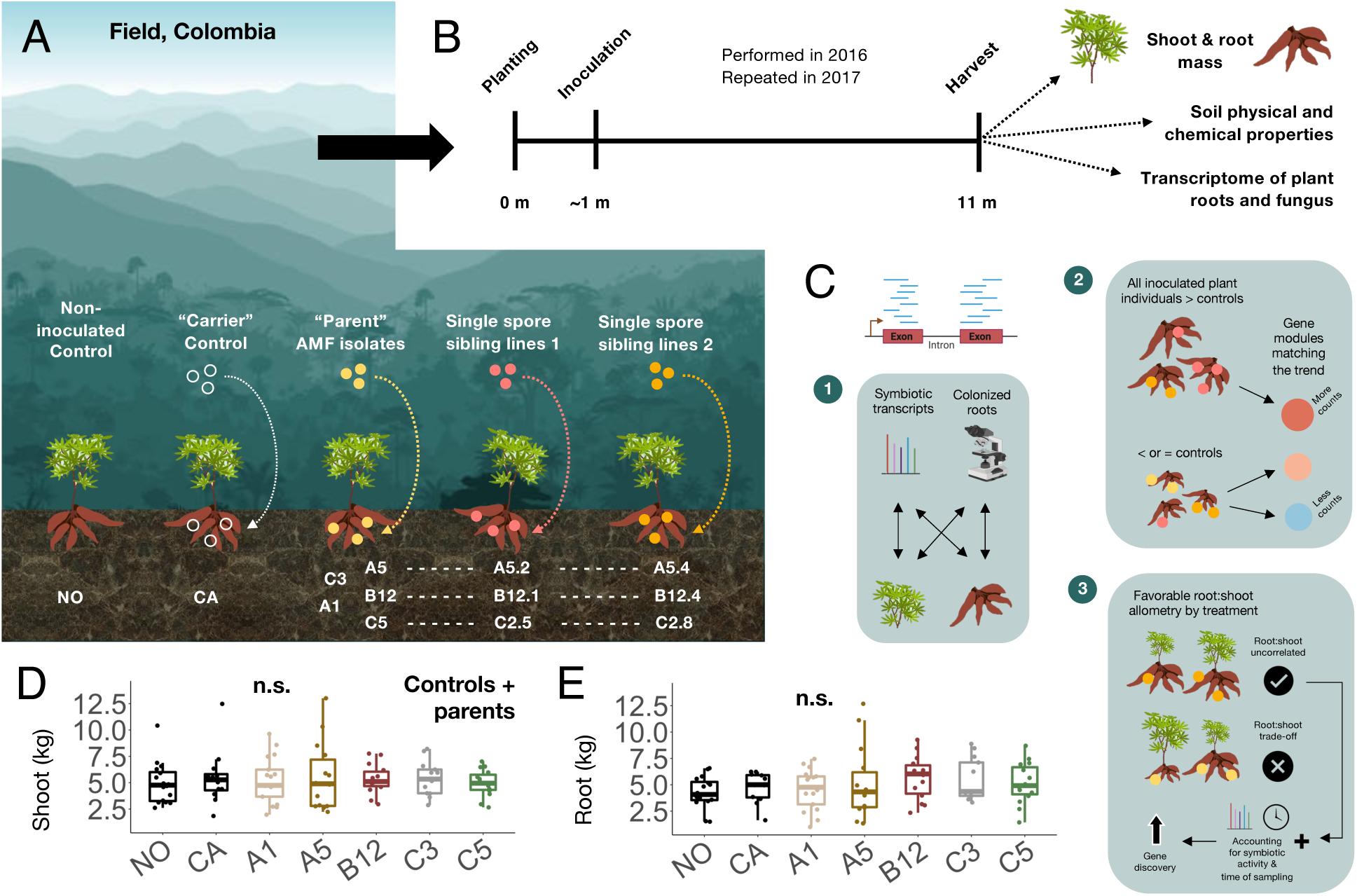
(A) Experimental treatments, (B) timelines and samples, (C) analysis workflows related to transcriptome data collected, (D) shoot and (E) root biomasses of the parental isolates. The latter two are presented as boxplots with medians (in kgs, y-axis) of the control or parental isolate treatments (x-axis) displayed. **n.s.**, not significant. Icons and images were generated using PowerPoint and BioRender.

The studied allometric trajectories were formed by the relationship (i.e., correlation) between shoot to root biomass. Shifts in allometry were represented by a strengthening or weakening of the correlation between these two traits, or changes in its slope or intercept. Comparing allometric and transcriptional changes within AMF “families” - among parental isolates and their single-spore progeny - allowed an exploration into whether repeated shifts in cassava allometry in particular AMF families could have single transcriptional origins. Just as allometry is a known ecological method that we newly applied to help parse the results of AMF interactions, we also drew from circadian theory in order to enhance our ability to understand cassava responses to AMF in the field, where they can especially be muddled by other experimental factors. The time each sample was taken in the field was determined from known plant and AMF core circadian gene transcription, and with this as well as our plant transcriptional symbiotic index (rather than colonization percentage) used as influencing factors in analyzing our transcription results, we were able to resolve more down- and up-regulated processes in both organisms that could be related to globally-important cassava yields.

## MATERIALS & METHODS

### Biological material and growth conditions

CM 4574-7 is an improved cassava (*Manihot esculenta*) cultivar, of industrial use, previously shown to respond positively to arbuscular mycorrhizal fungi (AMF) inoculation (Ceballos *et al*., 2019). It was grown from stakes in natural field conditions near Tauramena, Colombia (4.959329, −72.573215), in two consecutive field experiments. The experiments were separated in space and time within the same overall field area so as to avoid interactions from the inoculation of the previous year (Fig. S1A). Before each experiment, the area was cleared, tilled (3x harrow plow), treated with a pre-emergent herbicide (Active ingredient Diuron, Invesa, Colombia), and after planting, it was fertilized (100 Kg ha-1 urea & 100 Kg ha-1 diammonium phosphates, DAP, Monomeros, Colombia; 106 Kg ha-1 potassium chloride, KCl, Yara, Norway). Fertilization was equally divided between an application at 40- and 60-days post planting (dpp). The first experiment ran from September 2016 to August 2017 and the second from June 2017 to May 2018, representing planting and harvest dates, respectively.

Plants were inoculated with AMF in the field two weeks after the first leaves emerged from lateral meristems of the cassava stem cuttings (∼1 month post planting). The inoculum used in both experiments comprised 1 g of calcified diatomite containing ∼1000 AMF spores g^-1^ (Symbiom, Czech Republic); the spores of only one *R. irregularis* isolate was added to the base of each cassava individual. Eleven isolates, common to both the first and second experiments, were used: five (A1, A5, B12, C3, C5) originated from single spores from a field in Tänikon, Switzerland (Koch *et al*., 2004), subsequently referred to as the parental isolates, and two single spore sibling cultures originating as offspring from each of A5, B12 and C2, respectively. The two C2 offspring were considered like C5 offspring as the C2 and C5 parental isolates are genetically indistinguishable (Robbins *et al*., 2021). A detailed summary of the *R. irregularis* isolates used and previous characterizations can be found in Table S1.

Both field experiments had a randomized, blocked design. Each block contained one cassava individual from each inoculation treatment group, isolated by a physical barrier of nine non-inoculated plants from their next, differently-inoculated neighbor (Fig. S1B). Two control treatments were also included within each block: non-inoculated controls (“NO”) with nothing added at the time of inoculation, and carrier controls (“CA”) with 1 g of spore-free calcified diatomite powder added at the time of inoculation.

### Field measurements and sampling

At eight months post planting of the second experiment, samples of fine, fibrous roots from three random plants per treatment were collected from within a 10-15 cm circumference around the bases of inoculated cassava plants, at an approximate depth of 5-10 cm. These were sampled with minimal disturbance. Root samples were immediately flash-frozen in the field using a dry liquid nitrogen container (Cryoshipper, MVE Biological Solutions, USA) and kept below −80°C until RNA extraction for the purpose of generating an RNAseq dataset.

At harvest, fresh shoot and root biomass were separately weighed with a tared, digital, hand-held scale. Dried biomass was not measured, as it has previously been shown that for cassava, fresh and dry biomass values are tightly correlated (Ceballos *et al*., 2013, 2019; Peña *et al*., 2020). Additionally, fine fibrous roots of select treatments were collected at harvest in the same manner as at eight months, though they were placed instead in root a storage solution (99% ethanol:60% acetic acid, 3:1, v:v) until staining for AMF colonization quantification. When the log of all fresh root and shoot biomasses were taken to plot correlation graphs used for the allometric analyses, all samples with log_10_(shoot biomass) values less than 7.5 were removed due to observations in the field that these plants were constrained by non-representative edge conditions of the experiment. As these outliers only occurred in one year of the experiment, this further supported their removal. We present statistics related to their removal in Fig. S2A, as well as allometric trajectories separated per year in Fig. S2B. As the trajectories did not differ per year, data collected from both years was pooled unless otherwise specified.

After the harvest of the first experiment and before the planting of the second experiment, non-intact (i.e., disturbed) soil samples were collected from V-shaped pits dug at 10-15 cm depth (Soil Survey Staff, 2014). Soil organic carbon (SOC) was measured by the Walkley-Black method (Walkley & Black, 1934). Bulk density was measured by pycnometer, soil texture by the Boyoucos method, and acidity through KCl extractions (IGAC, 2006). Other elemental contents (e.g., sulfur, S; phosphorus, P) were determined by atomic absorption (Mehlich, 1984) and Bray II (Fassbender & Igue, 1967). Base saturation, or the percentage of each major cation with respect to the cation exchange capacity, was reported as % content.

### Sheaffer blue ink staining and root length colonization

Fine roots in root storage solution were processed and stained with Sheaffer blue ink (745, Germany) as described by Wilkes et al. (Wilkes *et al*., 2020). Root length colonization percentage (RLC %) was calculated using a grid-line intersection method (Giovannetti & Mosse, 1980).

### RNAseq sample preparation and sequencing

Three replicates of flash-frozen root samples from each treatment were randomly selected for the RNAseq analysis. They were homogenized, without unfreezing, using a liquid-nitrogen-supplied tissue mill (Cryomill, Retsch, Germany; 30 s @ 25 Hz, 30 s @ 5 Hz, 30 s @ 25 Hz) and processed using a plant RNA extraction kit and accompanying robot (Maxwell 16 LEV Plant RNA Kit, Promega, Germany). 2 uL total RNA was used to measure RNA quality (Fragment Analyzer, Agilent, USA) and only samples with an RNA integrity number above 7 were multiplexed and pooled together in a single library (TruSeq Stranded mRNA kit, Illumina, USA). Briefly, mRNA was selected and adapters were ligated using the Sciclone G3 Automated Liquid Handling Workstation (PerkinElmer, USA). Lane bias was accounted for by performing paired end sequencing of the same library across multiple lanes (Illumina HiSeq 4000, Illumina, USA). Approximately 50 million reads per sample were generated.

### RNAseq raw read processing

Raw, demultiplexed reads were trimmed using *trimmomatic 0.36* (*LEADING:28, TRAILING:28, SLIDINGWINDOW:10:28, MINLEN:50*). The ultimate goal was to quantify both cassava and *R. irregularis* reads. Traditional approaches map reads in a sequential manner using two separate genomes. This can lead to a disproportionate number of reads being misallocated to the first genome depending on conserved portions of genes across genomes (Espindula *et al*., 2019). Here, the reference genomes of *M. esculenta* (Phytozome, JGI: *Manihot esculenta* v7.1) and *R. irregularis* (Bioproject: PRJNA208392) were merged into a single reference genome to improve the process of dual-genome read mapping (Espindula *et al*., 2019). Reads were then mapped to the merged genome using *STAR 2.6.0c* with standard settings, on five threads.

Cassava and AMF read count data tables with only uniquely mapped reads were generated with *rsem 1.3.0*. The cassava and AMF datasets were separated for further processing due to large differences in detected plant and fungal reads. A per-sample library size normalization (TMM) was used to generate final counts (package: *edgeR,* function: *calcNormFactors* on a DGEList generated from the raw counts through *DGEList*) and enable comparison among samples (Robinson *et al*., 2010). TMM is more conservative than other normalizations, leading to fewer false positives but a lower detection of significantly differentially expressed (DE) genes (Dillies *et al*., 2013; Evans *et al*., 2018). Before differential analysis, TMM-normalized counts were filtered with *edgeR::filterByExpr* in order to ensure normal distributions for the plant and fungal datasets. 28113 plant genes and 2594 fungal genes were retained (of 33849 plant and 26143 fungal genes used for the mapping; 83.1% retained for the plant, 10.0% for the fungus). Total numbers of reads per treatment were checked after gene filtering and did not differ significantly (Fig. S3A, B). Total genes also did not differ significantly among treatments in either set (Fig. S3C, D).

### Retroactive determination of sampling time from RNAseq data

Plant circadian clock genes are well characterized and are known to peak at specific times of the day (Nakamichi, 2020). Using count abundances mapped to cassava’s “morning” (*MeLHY*) and “evening” (*MeTOC1*; Table S2) clock genes, with the knowledge that these genes negatively regulate each other (Gendron *et al*., 2012), two-hour ranges of the day were assigned to samples, representing the most likely times within which the sample was collected. This time was refined by the consultation of the count abundance of two additional circadian clock genes that peak between *MeLHY* and *MeTOC1*: namely, *MePRR5* and *MePRR7* (Fig. S4A).

*M. esculenta* clock gene identifiers used in the assignment of time of sampling are reported in Table S2. The estimated time of sampling was represented by the end of the range predicted (predicted range: 9-11, designation in the metadata column Sampling Time: 11, Table S3A). As a further, plant-independent confirmation of the assigned time of sampling, the count abundance of the *R. irregularis* clock gene *RiFRQ* (Table S2), described in previous literature as peaking in the morning hours (Lee *et al*., 2018), was plotted per assigned time of sampling and indeed peaked as previously described (Fig. S4B).

### Index of symbiotic transcriptional activity (ISTA)

Five cassava genes previously characterized to have ubiquitously increased transcript abundance among different cassava and AMF genotype combinations, and known to be absent when cassava is not inoculated (Mateus *et al*., 2019; Table S4), exhibited abundant counts in the normalized and filtered RNAseq data. Count numbers of these genes, however, differed in their degree of abundance and thus each had to be normalized to their presence in the first sample of the dataset in order to be compared. The average of the five normalized read counts in each sample was taken to represent that sample’s state of plant-AMF symbiotic activity, hereto referred to as its index of symbiotic transcriptional activity (ISTA).

Continuous values of the ISTA were used in determining correlations with shoot and root biomass values (metadata column ISTA-continuous, Table S3B). However, random effects included in linear mixed models are generally not continuous, therefore ISTA continuous values were also transformed into a categorical variable. Values between 0 and 0.075 were assigned as “LOW”, from 0.075 to 2.0 were “MED” (medium), and above 2.0 were “HIGH” ISTA values (metadata column ISTA-categorical, Table S3C).

### Root biomass rankings and CEMiTool analysis

Shoot and root cassava biomass, regardless of inoculation treatment, exhibited extensive variation (at minimum, 2.5-7.5 kg, max., 2.5-12.5 kg). The non-inoculated (NO and CA) cassava had lower maximum biomasses than most of the inoculated plants. We performed a treatment-independent analysis of plant transcription to find modules reflecting the higher root biomass maximums in inoculated plants. To do so, all plants were divided by ranks of root biomass production rather than treatment: plants with root biomasses above the maximum production of NO or CA plants were considered rank 1. Ranks 2-4 represented plants with a range of root biomasses that overlapped those of NO and CA plants; rank 4 plants had the lowest root biomasses among all plants. Divisions among ranks 2-4 were determined to produce a relatively even number of plants in each ranking group (Table S5; n = 7-8 plants).

CEMiTool was used to perform a gene set enrichment analysis by ranking group (version: 1.12.2; pipeline: Russo *et al*., 2018). Only the plant counts dataset was used, given that modules found to be significant from the CEMiTool pipeline were further explored by over-representation analysis, and this requires sufficiently resolved gene ontology assignments to the organism annotation. Most genes in the *R. irregularis* annotation are uncharacterized proteins.

### Statistical analyses

Statistical considerations for the CEMiTool analysis are described previously (Russo *et al*., 2018). Significances for boxplots are multiple comparisons extracted after significant ANOVA results. The data were fit to linear models and were graphically confirmed to fit the assumptions of normality and homoscedasticity, prior to ANOVAs and multiple comparisons (package: *emmeans*, version 1.7.0; Searle *et al*., 1980). This is also applicable to the correlation analyses, with the exception that *lstrends* (package: *emmeans*) was used to extract multiple comparisons, and Pearson’s correlations and significance are reported.

### RNAseq mixed modelling and downstream analysis

The plant and fungal RNAseq datasets were processed individually to find DE genes with unique expression patterns in B12.1- and C2.5-treated samples. Two different analyses were conducted: one without, and one with Sampling Time and ISTA included as random effects. Both involved fitting the count data to a linear mixed effects model with the fixed effect of the AMF treatment, with or without the random effects included (package: *voomWithDreamWeights*, function: *variancePartition*). Subsequently, DE statistics were calculated through the *eBayes* function (package: *limma*) and contrasts between treatments were extracted using *topTable* (package: *limma*). DE unique to C2.5 samples (found when C2.5 was compared to the other treatments of its family: C5 and C2.8), were extracted. The same was done for genes unique to the B12.1 treatment (in comparison to B12 and B12.4). Afterwards, only genes in common between these two extractions were kept. As an additional control, all genes with DE in contrasts between parental isolate treatments were excluded from the final set, as the phenotypes of interest did not manifest among these parental isolates. Heatmaps of results from the analysis without the random effects included and with them were generated using the *heatmap* function (package: *stats*).

To justify the inclusion of Sampling Time and ISTA as random effects, a variance decomposition analysis was performed to evaluate how much of the variance in gene count abundances was explained by these two variables versus by the AMF treatment. The model was re-fit with AMF treatment also as a random effect to produce an equivalent comparison of this treatment to the other phenotypes, and the function *fitExtractVarPartModel* was used (package: *variancePartition*). Percentages of variation explained by each tested random factor, as well as unexplained variation (in the residuals) was extracted for the plant and fungal gene counts and visualized as boxplots.

Finally, a weighted gene correlation network analysis (WGCNA) was run on samples treated with AMF isolates from the two families with the treatments of interest. The goal was to use genes resulting from the mixed model analysis (conservative) to pinpoint modules of interest within the WGCNA (less conservative), that could allow connections to be found among genes. The filtered, normalized and subset plant and fungal genes were used for the WGCNA analysis, which was the same input as for the mixed model analysis. Each set (plant vs. fungal genes) were run through the *DESeqDataSetFromMatrix* and *varianceStabilizingTransformation* functions (package: *DESeq2*). A threshold was picked for the WGCNA analysis through the *pickSoftThreshold* function (package: *WGCNA*) and used as the power input to the *blockwiseModules* (package: *WGCNA*, maxBlocksize = 10000). Resulting modules were searched for genes discovered through the mixed model analysis and modules with genes of interest were explored for significant correlations with other modules. Gene abundances in seven modules of interest were plotted against Sampling Time and ISTA as a final analysis of the influence of these random effects (Fig. S5).

## RESULTS

### Index of symbiotic activity correlates with shoot biomass and the trajectory differs by AMF inoculation treatment

There were no significant differences among the means of shoot (Fig. 1D) or root (Fig. 1E) biomass of cassava plants with and without inoculation with parental isolates. This appeared to result from the large variation in plant biomass in all treatments, including in the non-inoculated controls. Soil properties did not correlate with any cassava biomass results (Fig. S6).

Of the subset of plants with root colonization samples, there was no correlation between their biomass and root length colonization (RLC; Fig. 2A). We additionally used a plant index of symbiotic transcriptional activity (ISTA), developed in this study, to see if this correlated with cassava root or shoot biomass (Fig. 1C.1). Interestingly, the ISTA correlated significantly with cassava shoot biomass (Fig. 2B, upper panel). The direction of this correlation differed according to the inoculum treatment (Fig. 2B, significant ISTA:Treatment interaction). While shoot biomass in C3 inoculated plants decreased with an increasing ISTA, it increased for A1 and A5 inoculated cassavas, and neither increased nor decreased in the other treatments. There was no correlation between the ISTA and root biomass (Fig. 2B, lower panel). ISTA could not be directly compared to RLC as the subset of samples taken for colonization overlapped with only few samples randomly selected for RNAseq.

**Figure 2.**
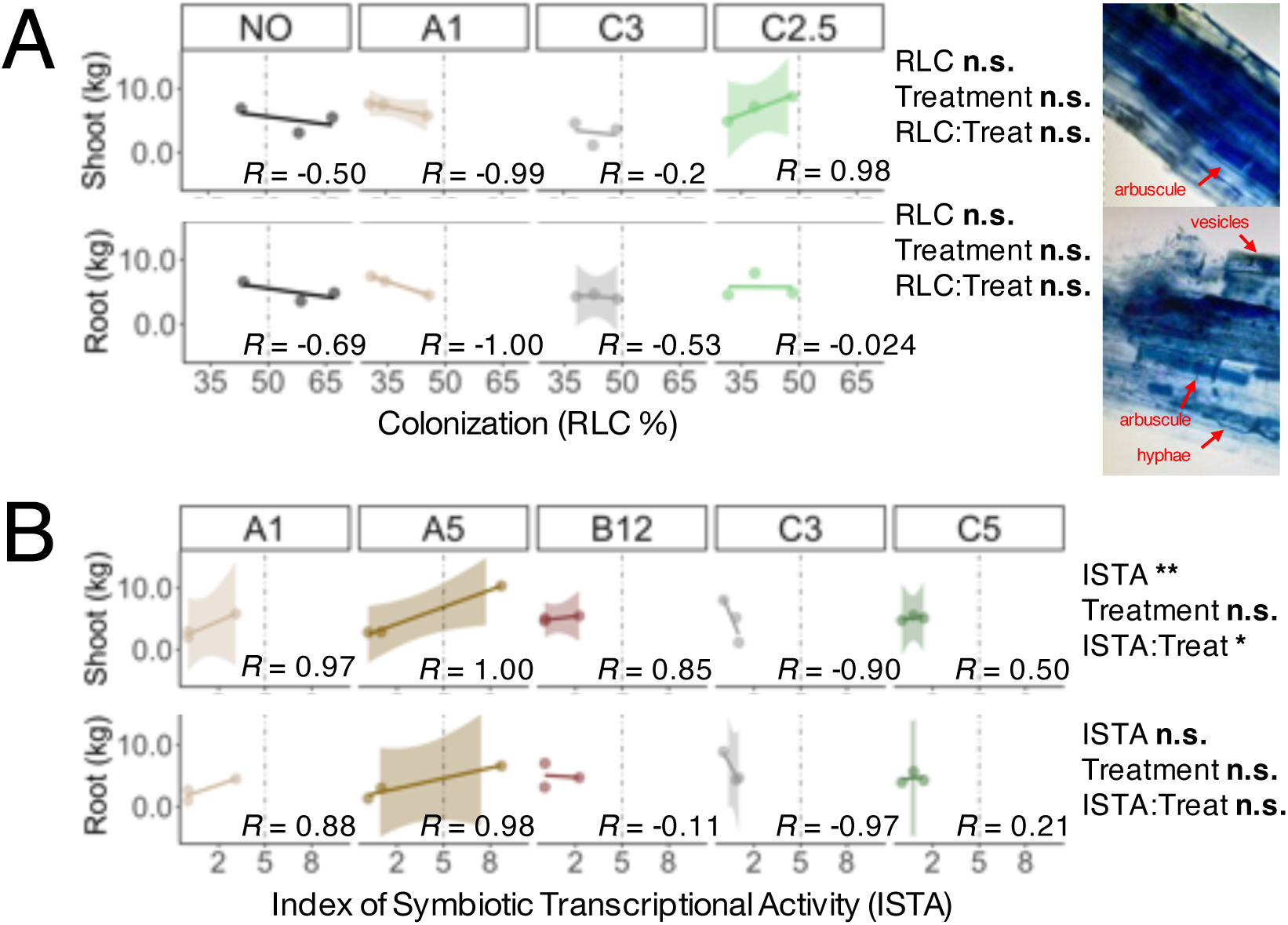
Correlation of (A) root length colonization percentage (RLC %, x-axis), as well as (B) the index of symbiotic transcriptional activity (ISTA, x-axis) with shoot and root biomasses (y-axes) of select plants within particular treatments (facets) from the experiment. ANOVA results are displayed to the right of each figure. **n.s.**, not significant; *, *p* value < 0.05; **, *p* value < 0.01. An inset displaying photos of the staining method used for the RLC % quantification is included to the far right of (A). Microscopic images by E.M.

### Intra-AMF family comparisons reveal significances among cassavas root biomass

Comparing inoculation treatments within their families (parental isolates and their single-spore progenies), allowed us to detect significant differences in mean cassava root biomass (Fig. 3A). However, mean cassava root biomass still only differed significantly among treatments within the families of the parental isolates B12 and C5. B12.1 inoculated plants exhibited significantly lower root biomass compared to the parental B12 treatment. None of the means of the C2 single-spore progeny differed significantly from those of the parental isolate C5, but the C2.8 isolate induced significantly lower cassava root biomass in comparison to the C2.5 treatment.

**Figure 3.**
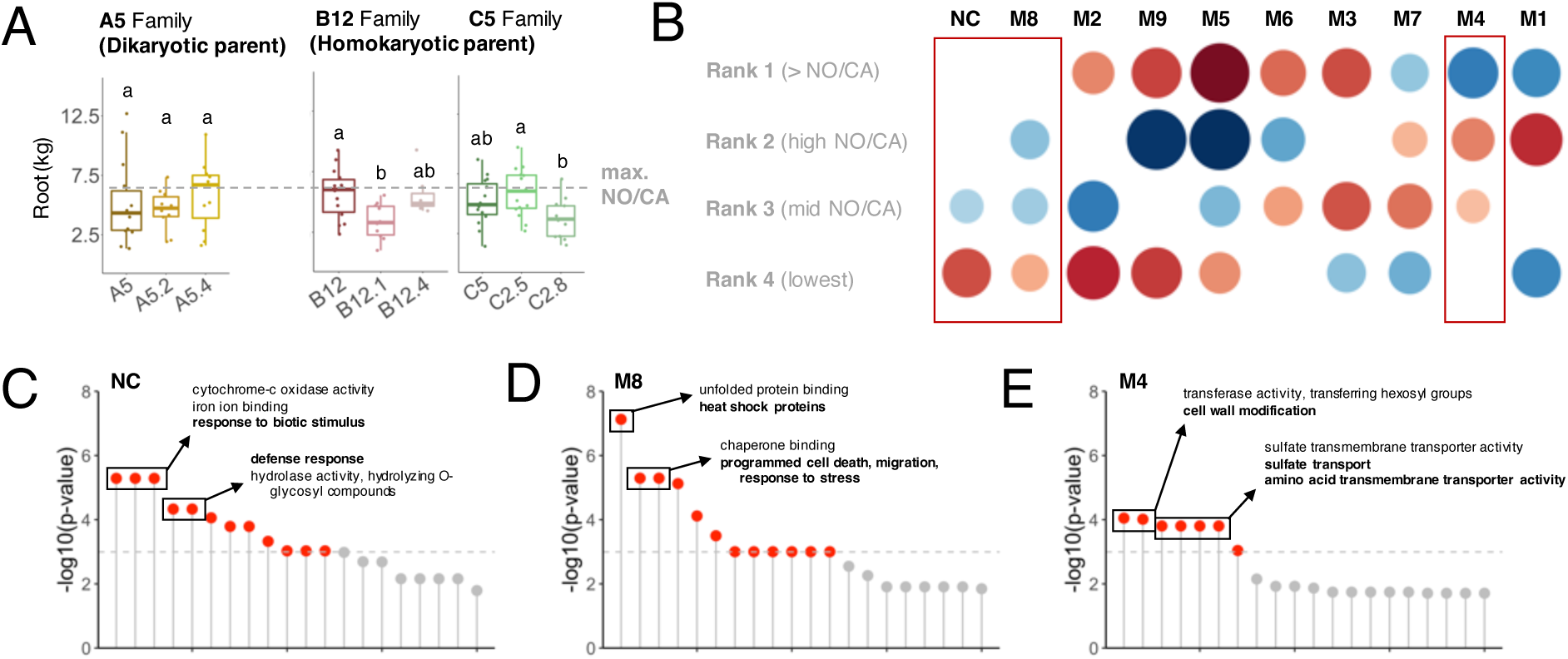
(A) Boxplot of median root biomass values (y-axis) for each treatment in the A5, B12 and C5 family isolates (x-axis). A dashed line across all facets represents the maximum root biomass observed in the control plants (treated with either no inoculum, NO, or with only the spore carrier powder but no spores, CA). Multiple comparisons are extracted from biomass means. Differing letters indicate significance among treatments where the *p* value < 0.05. (B) A CEMiTool analysis on plant genes, independent of treatment. All inoculated plants with RNAseq data were ranked by their root biomass production: rank one plant root biomasses exceed the maximum value of NO/CA plants, rank two through four represent descending values within the NO/CA biomass range. Red boxes mark modules with gene abundances changing continuously across ranks. Over-representation plots of plant genes in the modules of interest (genes on the x-axis, *p* value of their proportional representation on the y-axis, dashed line marking the significance threshold) are shown in (C) for the non-correlated, NC, module, (D) for M8 and (E) for M4. Gene ontologies of particular significance are indicated.

Though significant within-AMF family treatment effects generally involved a reduction in cassava root biomass, it was observed that all of the overall highest root biomass values occurred in inoculated plants (Fig. 3A): the root biomass maximum of nearly every inoculated treatment (10/11; Fig. 1E, 3A) was higher than the maximum values achieved by the non-inoculated, NO, or carrier powder only, CA, plants. This motivated our investigation into transcriptional changes, in common among AMF inoculations, that may lead to root biomass maximums (Fig. 1C.2).

### Root biomass maximums across treatments correlate with reduced transcriptional activity of plant stress genes

Cassava plants that contributed to the RNAseq dataset were assigned to ranks by their root biomass, regardless of the inoculum they received. A CEMiTool analysis found nine enriched plant gene sets – modules – that showed altered abundances across rank groups, though not necessarily in unison with the continuous decrease in root biomass (Fig. 3B). Modules with plant transcriptional patterns that followed a uniform decrease or increase across the rank groups, in conjunction with the changes in root biomass through the ranks, included only two modules (M8, M4), and the non-correlated module (NC), fitted this specification.

Surprisingly, all three of these modules of interest showed neutral or lower gene abundance in rank one plants (higher biomass). An over-representation analysis of the gene ontology (GO) terms associated to the plant genes in these three modules revealed genes for response to biotic stimulus and defense in the non-correlated module (Fig. 3C), heat shock and response to stress in M8 (Fig. 3D), and cell wall modifications, sulfate transport, and amino acid transport activity in M4 (Fig. 3E). Generally speaking, this indicated a lower investment in plant stress transcription (responses to biotic, abiotic stress) correlating to larger root biomass production that exceeded what NO and CA plants were able to achieve (details of genes per module: Table S6).

### A cassava tissue partitioning analysis reveals shifts towards favorable root production

Cassava root biomass plotted against shoot biomass (both logged for comparison on the same scale) revealed a weak allometric trajectory (i.e., significant correlation) between these two traits in control NO and CA cassavas (Fig. 4A, left). All parental isolate inoculations showed a strengthened relationship (i.e., these two variables were more closely correlated). In other words, shoot:root partitioning in cassava plants became more directed with the *R. irregularis* inoculations. The slope of the correlation did not differ among parental isolate treatments (i.e., no interaction was detected; Fig. 4A, right, and inset).

**Figure 4.**
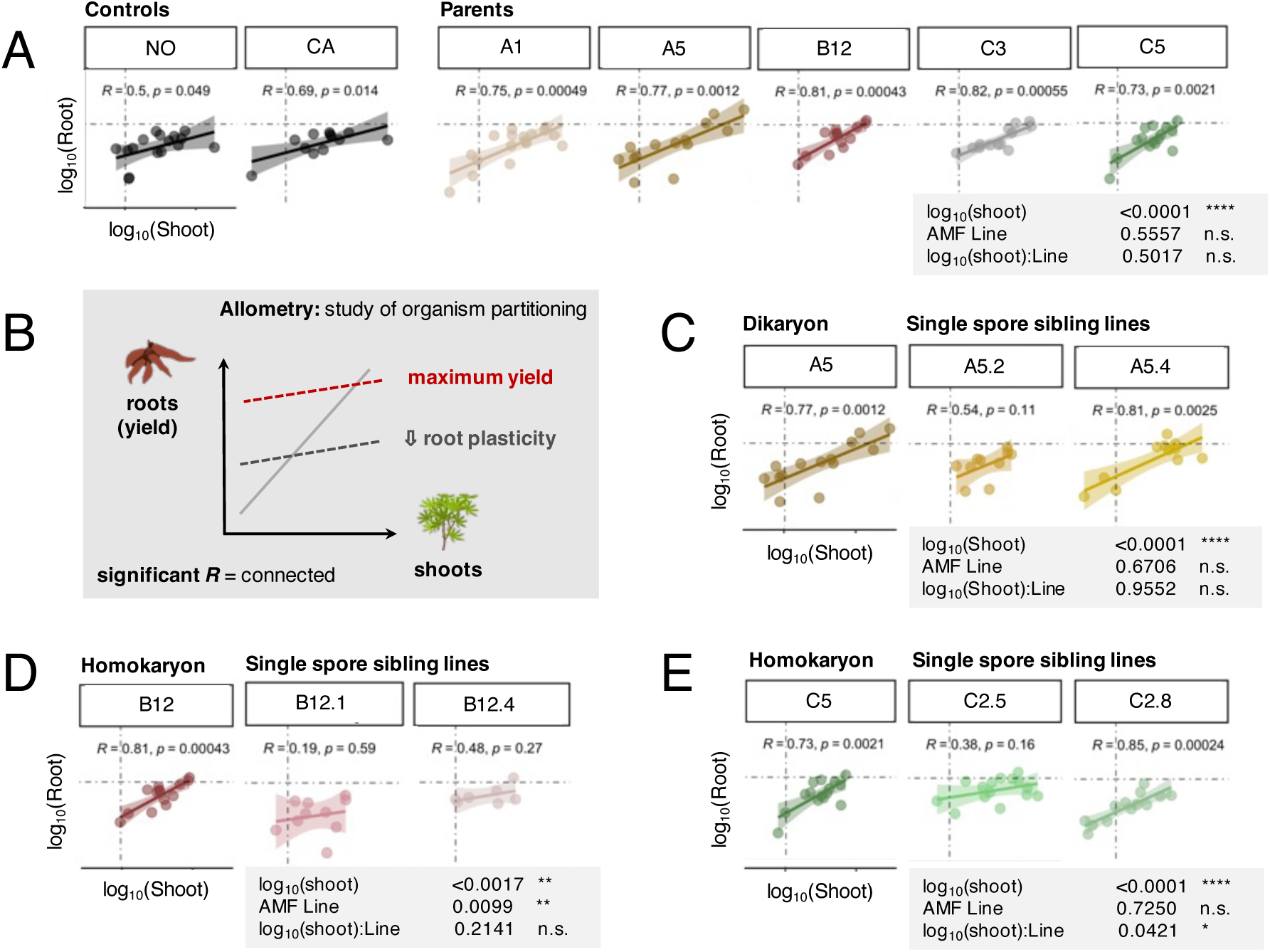
Allometric analyses of the shoot and root biomasses (log-scale, x- and y-axes, respectively) of cassava in (A) control (NO, CA) vs. parental isolate treatments. (B) An interpretation schematic is included to understand differences in allometric trajectories seen when intra-family comparisons are made for the (C) A5 family, (D) B12 family and (E) C5 family). *R* squared values and their significances are reported per inoculum and ANOVA results are displayed in gray boxes next to each figure (A, C-E). **n.s.**, not significant; *, *p* value < 0.05; **, *p* value < 0.01; ***, *p* value < 0.001; ****, *p* value < 0.0001.

Without a significant correlation (*R* value), two traits are not considered to form an allometric trajectory (dashed lines, Fig. 4B). A strong trajectory (solid line, Fig. 4B) is not always ideal: breaking the correlation between two traits can mean one trait receives a more consistent investment from the plant, regardless of the value of the other trait. This may sometimes be considered a decrease in plasticity in one trait (Fig. 4B, dashed gray line vs. solid gray line). For cassava, shoot:root partitioning would ideally not follow a trajectory and would be shifted toward a higher intercept on the root biomass axis, in order to increase and produce stable yields that are not reliant on aboveground biomass (Fig. 4B, dashed red line).

Intra-AMF family allometric analyses revealed that four single spore AMF progeny, one from each of the A5 and C5 families (A5.2, C2.5), and two from the B12 family (B12.1, B12.4), no longer induced the shoot:root allometric trajectories that had been strengthened by their parental isolate (Fig. 4C-E). This was only significant in the B12 and C5 families (ANOVA results in Fig. 4C-E, insets). Correlation between shoot and root biomass of cassava inoculated with A5.2 was weakened compared to those inoculated with A5, but the slope of the potential trajectory was unchanged. An unaltered slope does not affect the extent of variation in the plant’s shoot and root biomass values (Fig. 4B, C).

Not only was the allometric shoot and root biomass trajectory absent in cassava inoculated with B12.1, but the root biomass values became more consistent, albeit at a lower production level (Fig. 4D). Alternatively, plants inoculated with B12.4 demonstrated the ideal removal of the shoot:root biomass trajectory with stabilized root production occurring at the maximum values of root biomass (Fig. 4D). A similar effect was also observed in cassava inoculated with single spore line C2.5 when compared to the effect of its other AMF family members (Fig. 4E). B12.4 and C2.5 thus became ideal treatments to perform further transcriptional analysis; cassava and *R. irregularis* gene abundances uniquely present in samples treated with these progenies could be targeted.

### Including ISTA and time of sampling as random effects in a differential expression (DE) analysis improved discovery of AMF-induced down-regulated genes

A pipeline to extract significantly DE genes unique to B12.4- and C2.5-inoculated cassava plants produced 21 up-regulated plant genes and 1 up-regulated fungal gene (Fig. 5A, B). However, both plant and fungal gene counts were significantly influenced by the ISTA of their sample of origin, as well as by the time of day that the sample was collected (retroactively determined through circadian clock gene abundances: Sampling Time). The amount of variance in gene abundances explained by the ISTA or Sampling Time of a sample generally exceeded that explained by the AMF treatment (Fig. 5C). This was especially true for plant gene abundances (Fig. 5C, left); for fungal genes, ISTA, Sampling Time, and AMF explained relatively equal portions of count variance (Fig. 5C, right). Including ISTA and Sampling Time as random effects in the DE genes pipeline led to a 268% increase in DE gene identification: 45 up-regulated plant genes, 12 down-regulated plant genes, and 2 up-regulated fungal genes (Table 1; Fig. 5D). Only with the inclusion of these random effects were down-regulated plant genes found related to the allometric phenotypes of interest.

**Figure 5.**
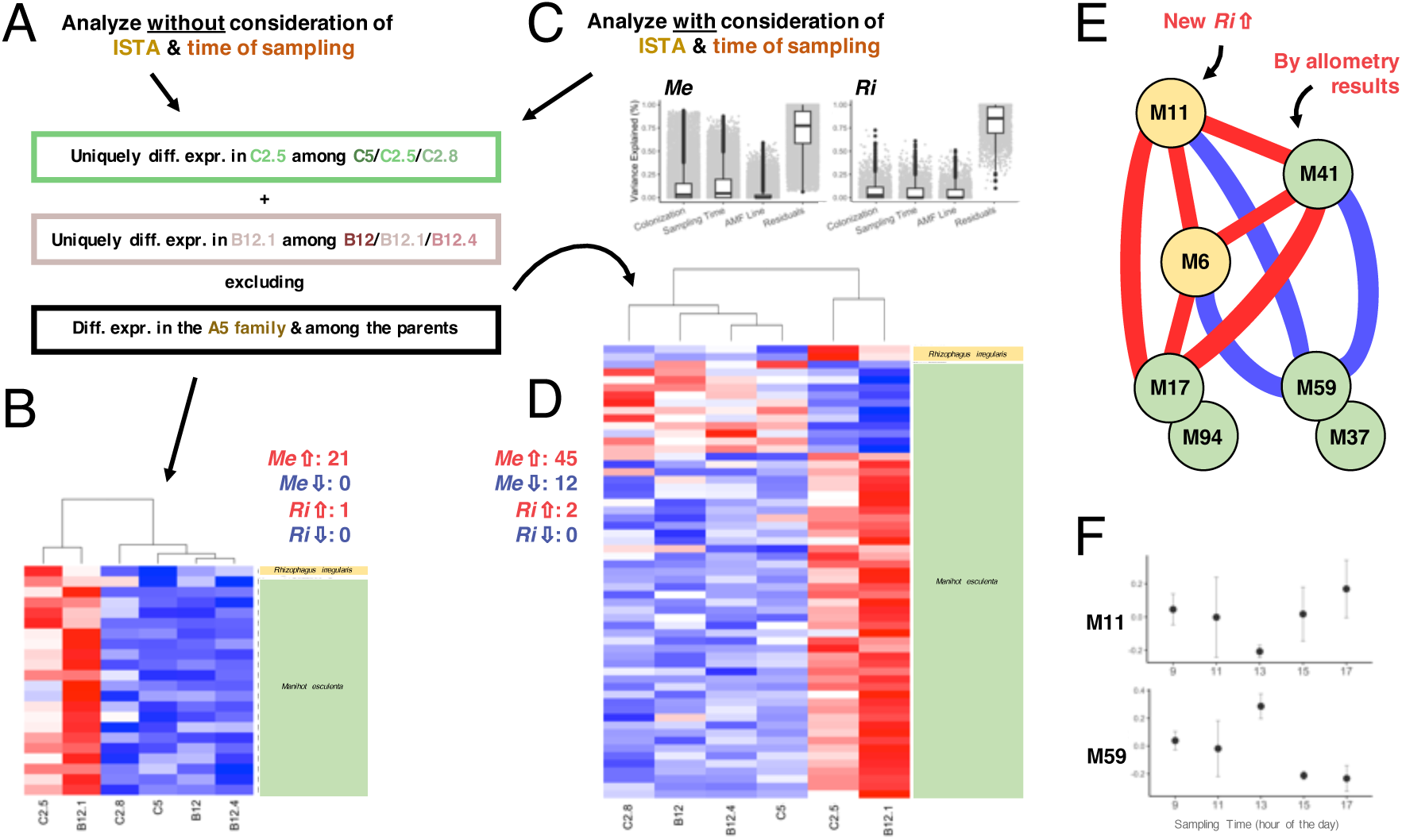
(A) The pipeline for the differential expression (DE) analysis performed based on the allometric results, either without (B) or with (C, D) acknowledgement of the random effects of the index of symbiotic transcriptional activity (ISTA) and time of sampling (Sampling Time). (B) Relative gene transcript abundances across the C5 and B12 families of significant genes found without accounting for random effects; significant plant genes are shown on the bottom (green, *Me*) and fungal genes on the top (yellow, *Ri*). (C) Boxplots of the median variance explained per gene (%, y-axis) by Colonization, Sampling Time, the AMF, and the residuals (x-axis), for *Me* (left) and *Ri* (right) genes, justifying the inclusion of these random effects in a second model. This second analysis produced (D) an extended list of DE genes. (E) Visualization of the results of a weighted gene correlation network analysis (WGCNA) informed by the DE findings when random effects were included. (F) The eigenvalues (y-axis, ± SE) of two modules of interest from the WGCNA plotted against Sampling Time (x-axis).

**Table 1.**
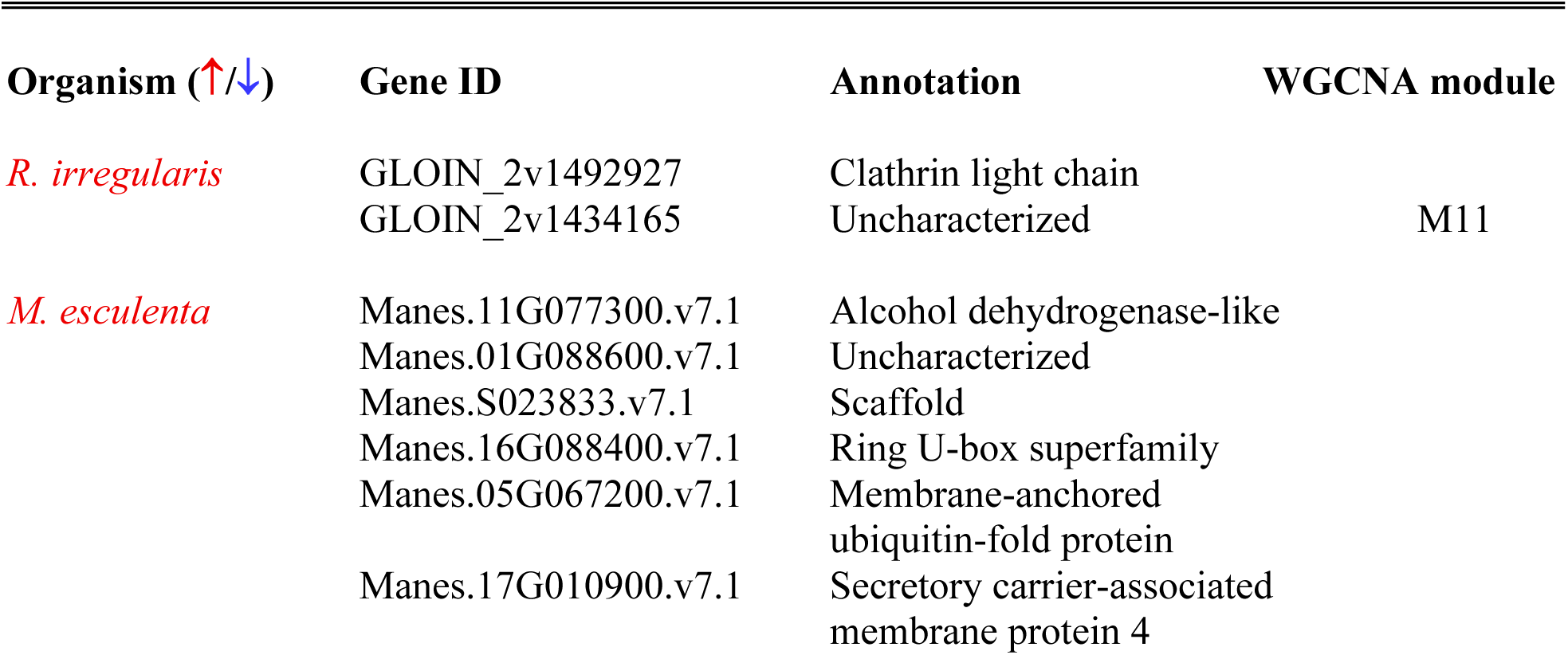
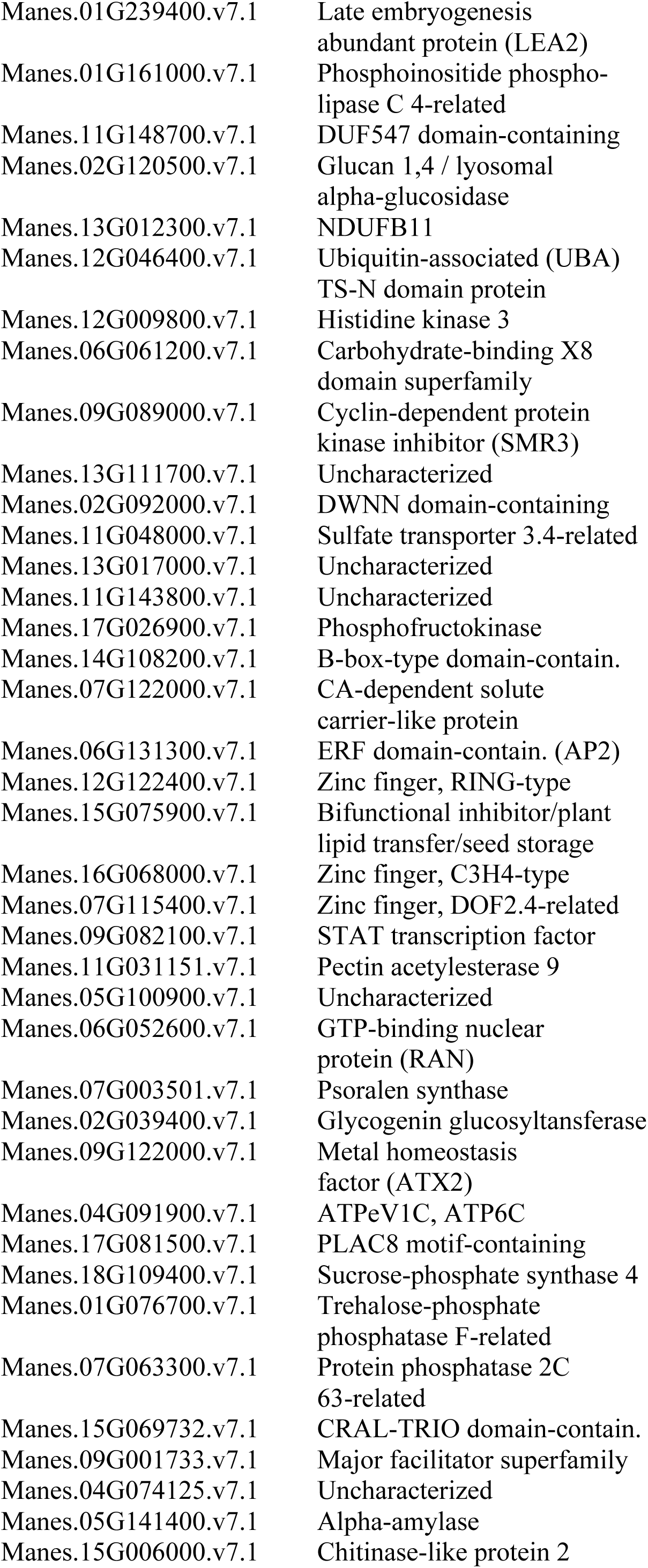
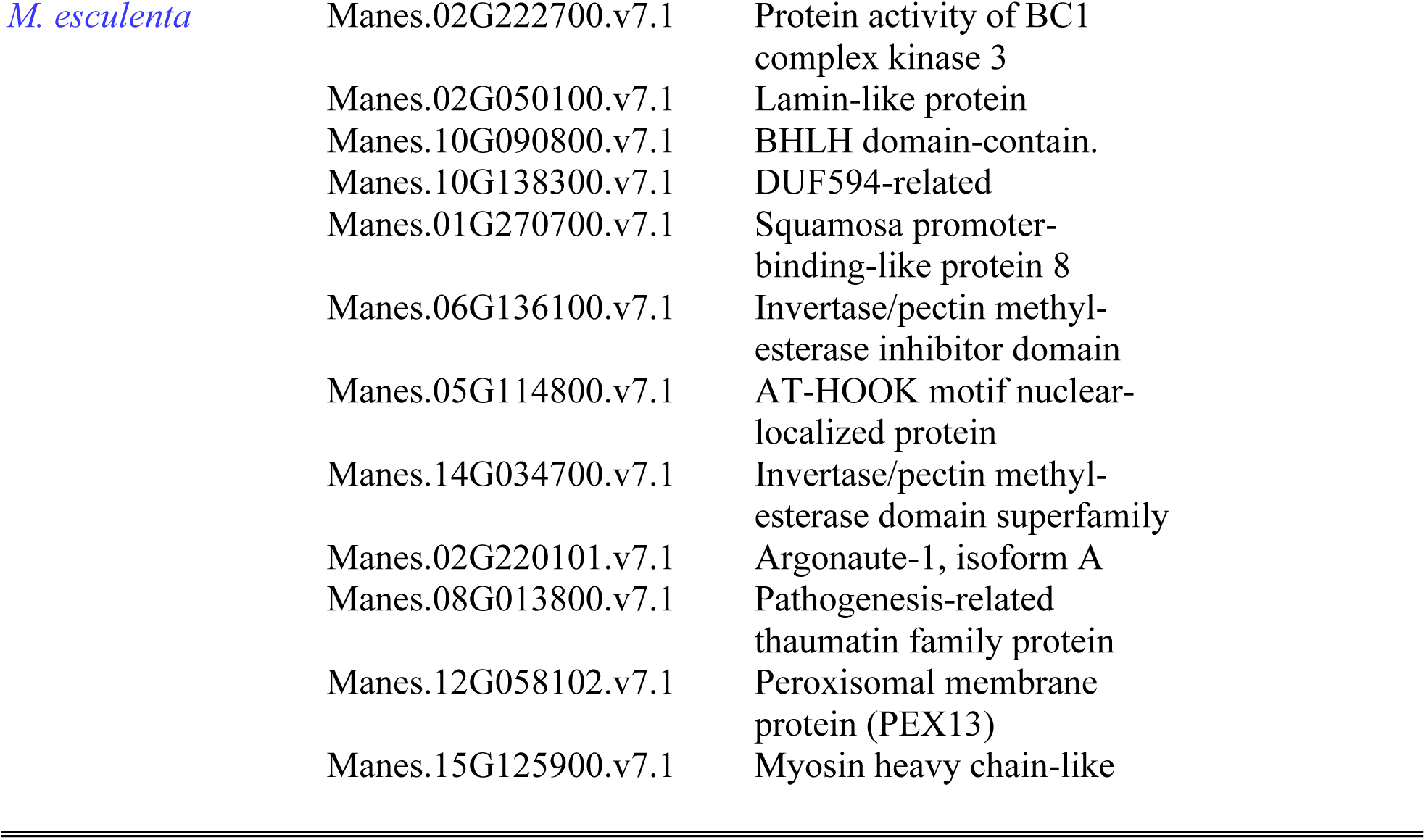
Genes of interest (IDs, annotations) identified from the DE analysis and their locations in the modules of interest produced from the WGCNA.

### A network analysis finds correlated plant and fungal modules supported by the DE results

Weighted gene correlation network analyses (WGCNA) run using only B12 and C5 family samples detected four significant plant modules (out of 115 modules detected), but no significant fungal modules (out of 22). Only one of the four significant plant modules, M41, revealed gene count abundance patterns that were unique to B12.4 and C2.5 cassava roots. The inability to detect a significant module in the fungal count dataset was relieved by the previous DE results, where the newly detected *R. irregularis* gene found in the second DE analysis (with random effects included) was able to be located in one of the WGCNA fungal modules: M11. The M41 plant module and M11 fungal module were highly positively correlated (Fig. 5E).

Only one fungal module, M6, was significantly correlated to both the fungal M11 and plant M41, forming a tri-partite positive correlation (Fig. 5E). To reduce the number of additional plant modules correlated to the plant M41 and fungal M11, they were filtered to only those also correlating with the fungal M6, resulting in four additional plant modules of interest: plant M17 and M94 positively correlated with, and plant M59 and M37 negatively correlated with the plant M41 and fungal M11 and M6 modules (Fig. 5E). WGCNA module locations of genes found in the DE analysis are reported in Table 1 (all genes in significant modules: Table S7).

### Plant and fungal modules of interest follow diurnal rhythms

Given the influence of the ISTA and Sampling Time on plant and fungal counts, the eigenvalues of the final WGCNA modules of interest were plotted against the ISTA categories as well as the Sampling Time points. No clear connection was found from categorizing the module expressions by ISTA, but all modules show clear diurnal patterns of expression (fungal example: Fig. 5F, upper; plant example: Fig. 5F, lower; all: Fig. S5).

## DISCUSSION

We introduce a new methodology using input from several different disciplines and increase both gene discovery and our understanding of *R. irregularis* effects on cassava production after inoculation in the field. Specifically, we analyzed in-field *R. irregularis* transcription, and cassava transcription, and link this to cassava biomass partitioning (Fig. 1). A plant index of symbiotic transcriptional activity (ISTA), aiming to represent plant differences in their active involvement in symbiosis with AMF, predicted shoot biomass in some treatments but never root biomass. Asking whether single-isolate treatments in general can increase cassava root biomass maximums, and what plant transcriptional processes may be linked to these increases, revealed down-regulated plant gene modules in common among cassavas that exceeded control plant root production. Only allometric analyses of cassava root and shoot biomasses and comparing treatments within isolate families allowed us to specify positive impacts from single-isolate inoculation on cassava outcomes. Differentially expressed (DE) genes in common to two isolate treatments, both that led cassava root production to be disconnected from its shoot biomass, included only plant genes with increased transcription in the roots of interest. After including ISTA and transcription sampling time as random effects, we were able to newly reveal down-regulated plant gene candidates, which was more in support with our previous analysis on cassava root biomass maximums.

Though the specific transcription of the inoculated isolate could not be tracked after application, all transcript abundances and yield changes were interpreted as downstream consequences of differences in the single-isolate inoculum. AMF transcription represents the total resulting *R. irregularis* transcription, including the inoculated isolate and the background *R. irregularis* community. It cannot be excluded that some AMF transcripts that we mapped to the *R. irregularis* model genome may have been from other AMF species in the original field soil. *R. irregularis* gene candidates should thus be approached with caution. It is to note though that the *R. irregularis* DE genes we found had transcription that was unique to the roots of cassavas with a given isolate treatment. Promising gene candidates should still be explored for their presence and transcription among and within AMF species to follow up on their potential role in *R. irregularis* and cassava interactions.

There is increasing emphasis on the fact that plant growth changes more likely link to the extent of transcriptional activity at the plant-AMF interface, rather than root length colonization (Hewins *et al*., 2015; Kobae, 2019). Indeed, that shoot biomass, but not the root biomass, of cassava plants correlated to plant’s ISTA introduces a novel understanding of where the effects may manifest of different plant responses to the AMF symbiosis. As AMF rely on carbon from their plant hosts (Shi *et al*., 2023), it is logical that investment in the AMF symbiosis would be positively correlated with the size of the photosynthesizing canopy. Surprisingly, our ISTA correlations revealed both positive and negative relationships between cassava shoot biomass and the extent of plant-AMF symbiotic activity. Clearly, symbiotic transcription can be highly altered, in several directions. The ISTA findings do need to be further replicated, but they do present an interesting future research avenue.

The allometric analyses were most effective in parsing field cassava biomass variation to then link it to single-isolate *R. irregularis* treatments. By plotting allometric trajectories showing the partitioning of each cassava plant between its shoot and root biomass, trends could be found in consistent partitioning regardless of the size of individual cassava plants within treatment, which was the main cause of the observed variability. This method thus takes advantage of variability, rather than suffering from it. If all the individual plants fell on a consistent line of partitioning, otherwise known as having a directed allometric trajectory, then we would observe a high and significant correlation among shoot and root biomass in that particular treatment. This work first reveals that control cassava in the field, with their background AMF communities, only exhibited weak allometric trajectories (Fig. 4). Cassava inoculated by the parental isolates all showed significantly strengthened trajectories, although the allometric trajectories formed by these were not different among these isolates. Regardless, this has important implications for the predictability of yields of cassava, given inoculation with only 1000 spores of each AMF isolate had a consistent effect on this. Intra-AMF family comparisons then produced variances in allometric trajectories that could be used to drive transcriptomic analyses.

This study reveals new *M. esculenta* and *R. irregularis* gene targets for the optimization of cassava tuber production through AMF interactions (Fig. 5; Table 1). The significant effect of including ISTA and sampling time as random factors, which led to the discovery of one of the *R. irregularis* genes and all of the down-regulated *M. esculenta* genes, underlines their importance in reducing noise in field transcriptomic samples. Most importantly, this work takes a crucial step forward in making field transcription analyses on a widespread symbiotic relationship more accessible. After the random effects reduced noise in the data, our ranking module analysis (Fig. 3), DE gene determination, and WGCNA approaches (Fig. 5) helped to co-validate gene candidates found through methods with different levels of conservativeness. These strategies, in addition to the use of allometric trajectories to guide the DE contrasts, can be used on existing as well as new field datasets to uncover more genes related to outcomes from the plant-AMF symbiosis under natural or agricultural conditions.

The time of sampling was retroactively determined for our field samples using gene transcription from core circadian clock genes in the plant. These are highly conserved across plants, and as large percentages of genes in plants are diurnally rhythmic, the importance of considering timing in plant gene transcription analyses has been previously emphasized (Ferrari *et al*., 2019). Given that *R. irregularis* also has a characterized circadian clock (Lee *et al*., 2018), the two partners in the plant-*Ri* symbiosis could interact at specific times in their circadian rhythms, contributing to the influence of this effect in our DE mixed models. Whether time of sampling is known in previous datasets or not, it can be determined through the method we present in our work. Abundance of cassava genes of interest followed the peaking times of the *R. irregularis* gene abundances, either in tandem (up-regulated) or in opposition (down-regulated). It is interesting that ISTA, and thus symbiotic involvement as recorded from the plant side of the interaction, may only match with the afternoon peaking timepoint of *R. irregularis* presence and transcription (Fig. S7). Investigating the role of each of these factors in altering allometric shifts of cassava plants under more controlled conditions would be the next step to verify these findings.

Finally, remaining open questions include whether the isolates of interest in this study can cause allometric shifts, as observed in the field, when they are inoculated on the same cassava variety in otherwise sterile soil in the glasshouse. This would indicate whether microbiome, soil, or other field-specific environmental factors may be the mechanism or feedback through which a single-spore AMF inoculum confers its effect. This set up could also evaluate the consistency of transcription of the single inoculated *R. irregularis* isolate with that of the totality of the *R. irregularis* community, which is what we were able to measure in our field cassava plant roots. This would build upon our advances, which showed, for the first time, that some single-spore inoculums of *R. irregularis* can strengthen cassava tissue partitioning in the field, some can cause allometric shifts towards stabilized yields regardless of shoot biomass, and that the latter effect is linked to specific, diurnal transcriptional patterns in cassava and in the AMF *R. irregularis*. Our study is a crucial step in determining the mechanisms by which plant-AMF symbioses can improve crop yields, in addition to providing new tools such as the ISTA and transcriptional study approaches for use in further plant-AMF field studies.

## ACKNOWLEDGEMENTS

We thank Mr. Lino Vega for the lease of the field sites in Tauramena, Colombia, and for providing tools and the cassava material used in the experiments. We thank the National University of Colombia for regulatory oversight and support of D.C.P-Q. and I.C., and the State of Vaud for their support of C.R. and E.M.

## COMPETING OR CONFLICTS OF INTEREST

The authors declare no conflicts of interest with this work.

## AUTHOR CONTRIBUTIONS

Conceptualization: C.R., I.R.S., D.C.P-Q., A.R.; Methodology: C.R., D.C.P-Q., I.C.; Investigation: C.R., D.C.P-Q.; Data Curation: C.R., D.C.P-Q., E.M.; Formal Analysis: C.R., E.M.; Interpretation: C.R., E.M., I.R.S.; Resources: I.R.S., A.R.; Writing - Original Draft: E.M.; Writing - Review and Editing: E.M., I.R.S., C.R., D.C.P-Q., A.R., I.C.; Visualization: E.M.; Supervision and administration: C.R., I.R.S.; Funding Acquisition: I.R.S.

## DATA AVAILABILITY

The raw, demultiplexed RNAseq read dataset generated during and/or analysed during the current study are available in the European Nucleotide Archive (accession: PRJEB44128).

## SUPPORTING INFORMATION

**Fig. S1** Schematics of the (A) field site and treatment blocks per year, and the (B) 9-plant design per treatment.

**Fig. S2** Graphical and statistical justification for (A) shoot outlier removals and (B) the combining of data from both field experiment years for the allometric analyses.

**Fig. S3** Analysis of (A, B) reads and (C, D) genes per treatment using normalized read count data for the plant (A, C) and fungus (B, D).

**Fig. S4** Circadian gene transcription (y-axis) plotted across time (x-axis) for the (A) plant and (B) fungus to validate sampling time assignment.

**Fig. S5** Eigenvalues of all of the final 7 WGCNA modules of interest plotted against (A) Index of Symbiotic Transcriptional Activity (ISTA), and (B) sampling time.

**Fig. S6** Soil properties correlated with (A) shoot and (B) root biomass results.

**Fig. S7** Putative peaking times of the (A) plant Index of Symbiotic Transcriptional Activity (ISTA) and (B) *R. irregularis* presence in plant roots, as determined by an *Ri* housekeeping gene.

**Table S1** *R. irregularis* isolates used and studies where they were previously characterized.

**Table S2** *M. esculenta* and *R. irregularis* clock gene IDs.

**Table S3** Metadata for the samples taken for transcriptomic analysis including retroactively determined sampling time, and Index of Symbiotic Transcriptional Activity (ISTA) as a continuous and categorical variable.

**Table S4** *M. esculenta* gene IDs used to calculate the Index of Symbiotic Transcriptional Activity (ISTA) raw (continuous) values.

**Table S5** Information on the ranking group divisions for the CEMiTool analysis.

**Table S6** Details on genes in the modules of interest from the CEMiTool analysis (NC, M8, M4). *Submitted as an Excel file*.

**Table S7** Details on genes in the final 7 WGCNA modules of interest. *Submitted as an Excel file*.

