## Supporting Information for "Diurnal plant and *Rhizophagus irregularis* transcriptional patterns are linked to shifts in cassava tissue partitioning in the field"

New Phytologist

***New Phytologist* Supporting Information**

Article title: Diurnal plant and *Rhizophagus irregularis* transcriptional patterns are linked to shifts in cassava tissue partitioning in the field
Authors: Erica McGale, Chanz Robbins, Diego Camilo Peña-Quemba, Isabel Cristina Ceballos, Alia Rodriguez, Ian R. Sanders
Article acceptance date: Click here to enter a date.

The following Supporting Information is available for this article:

**Fig. S1** Field site, treatment and design schematics.

**Fig. S2** Graphical and statistical justifications for allometric data combination and processing.

**Fig. S3** Reads and genes per treatment after normalization of read count data.

**Fig. S4** Circadian gene transcription plotted across time for sampling time assignment validation.

**Fig. S5** Eigenvalues of the WGCNA modules of interest plotted against the random factors.

**Fig. S6** Soil properties correlated with shoot and root biomass results.

**Fig. S7** Putative peaking times of plant symbiotic activity and *R. irregularis* presence in roots.

**Table S1** *R. irregularis* isolates used and studies where they were previously characterized.

**Table S2** *M. esculenta* and *R. irregularis* clock gene IDs.

**Table S3** Metadata for the samples taken for transcriptomic analysis.

**Table S4** *M. esculenta* gene IDs used for the Index of Symbiotic Transcriptional Activity.

**Table S5** Information on the ranking group divisions for the CEMiTool analysis.

**Table S6** Genes in modules of interest from the CEMiTool analysis. *Submitted as an Excel file.*

**Table S7** Genes in the final 7 WGCNA modules of interest. *Submitted as an Excel file.*

**Fig. S1** Schematics of the (A) field site and treatment blocks per year, and the (B) 9-plant design per treatment.

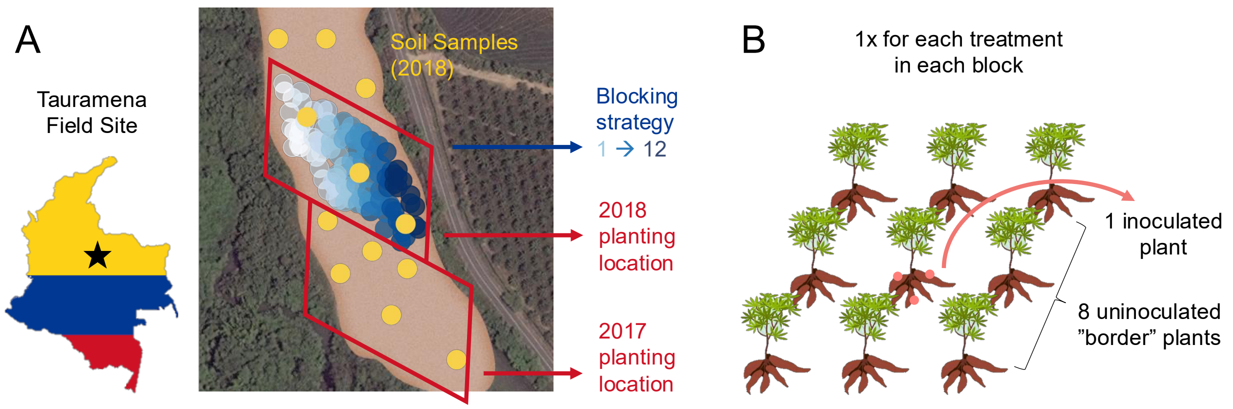

New Phytologist

**Fig. S2** Graphical and statistical justification for (A) shoot outlier removals and (B) the combining of data from both field experiment years for the allometric analyses.

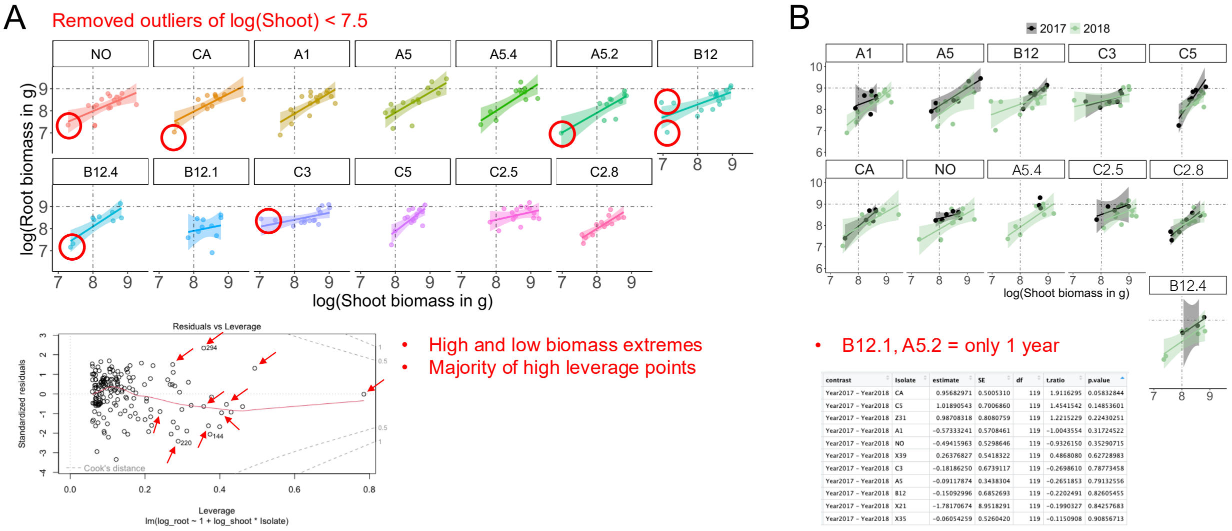


| contrast | Isolate | estimate | SE | df | t.ratio | p.value |
|---|---|---|---|---|---|---|
| Year2017 - Year2018 | CA | 0.95682971 | 0.5005310 | 119 | 1.9116295 | 0.05832844 |
| Year2017 - Year2018 | C5 | 1.01890543 | 0.7006860 | 119 | 1.4541542 | 0.14853601 |
| Year2017 - Year2018 | Z31 | 0.98708318 | 0.8080759 | 119 | 1.2215229 | 0.22430251 |
| Year2017 - Year2018 | A1 | -0.57333241 | 0.5708461 | 119 | -1.0043554 | 0.31724522 |
| Year2017 - Year2018 | NO | -0.49415963 | 0.5298646 | 119 | -0.9326150 | 0.35290715 |
| Year2017 - Year2018 | X39 | 0.26376827 | 0.5418322 | 119 | 0.4868080 | 0.62728983 |
| Year2017 - Year2018 | C3 | -0.18186250 | 0.6739117 | 119 | -0.2698610 | 0.78773458 |
| Year2017 - Year2018 | A5 | -0.09117874 | 0.3438304 | 119 | -0.2651853 | 0.79132556 |
| Year2017 - Year2018 | B12 | -0.15092996 | 0.6852693 | 119 | -0.2202491 | 0.82605455 |
| Year2017 - Year2018 | X21 | -1.78170674 | 8.9518291 | 119 | -0.1990327 | 0.84257683 |
| Year2017 - Year2018 | X35 | -0.06054259 | 0.5260420 | 119 | -0.1150908 | 0.90856713 |

**Fig. S3** Analysis of (A, B) reads and (C, D) genes per treatment using normalized read count data for the plant (A, C) and fungus (B, D).

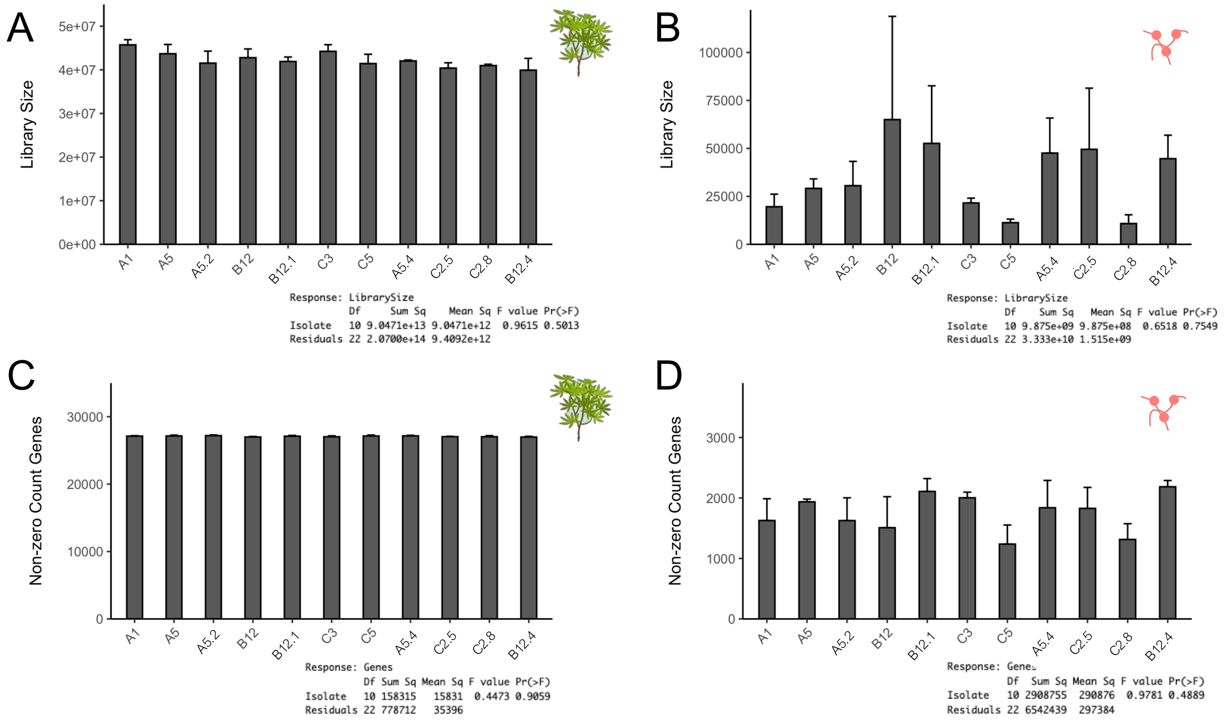

**Fig. S4** Circadian gene transcription (y-axis) plotted across time (x-axis) for the (A) plant and (B) fungus to validate sampling time assignment.

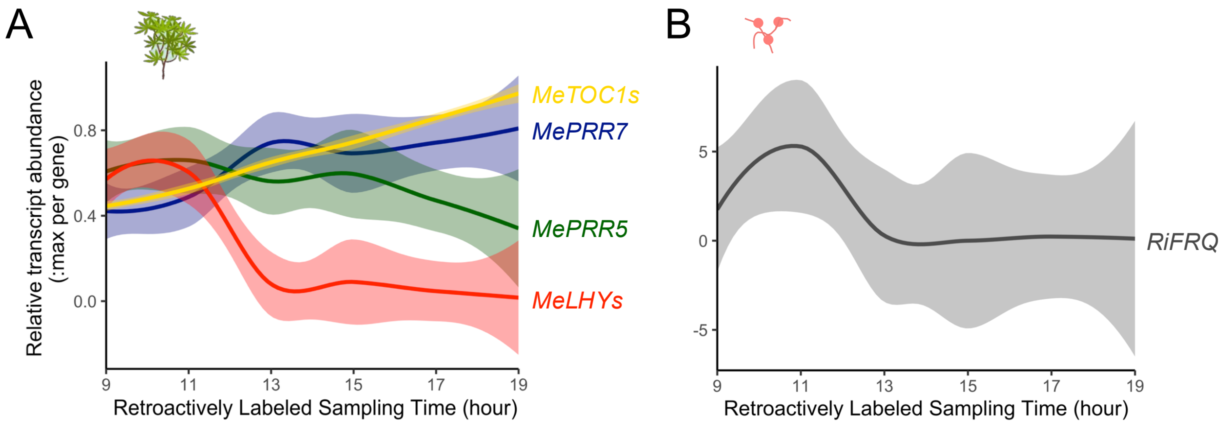

**Fig. S5** Eigenvalues of all of the final 7 WGCNA modules of interest plotted against (A) Index of Symbiotic Transcriptional Activity (ISTA), and (B) sampling time.

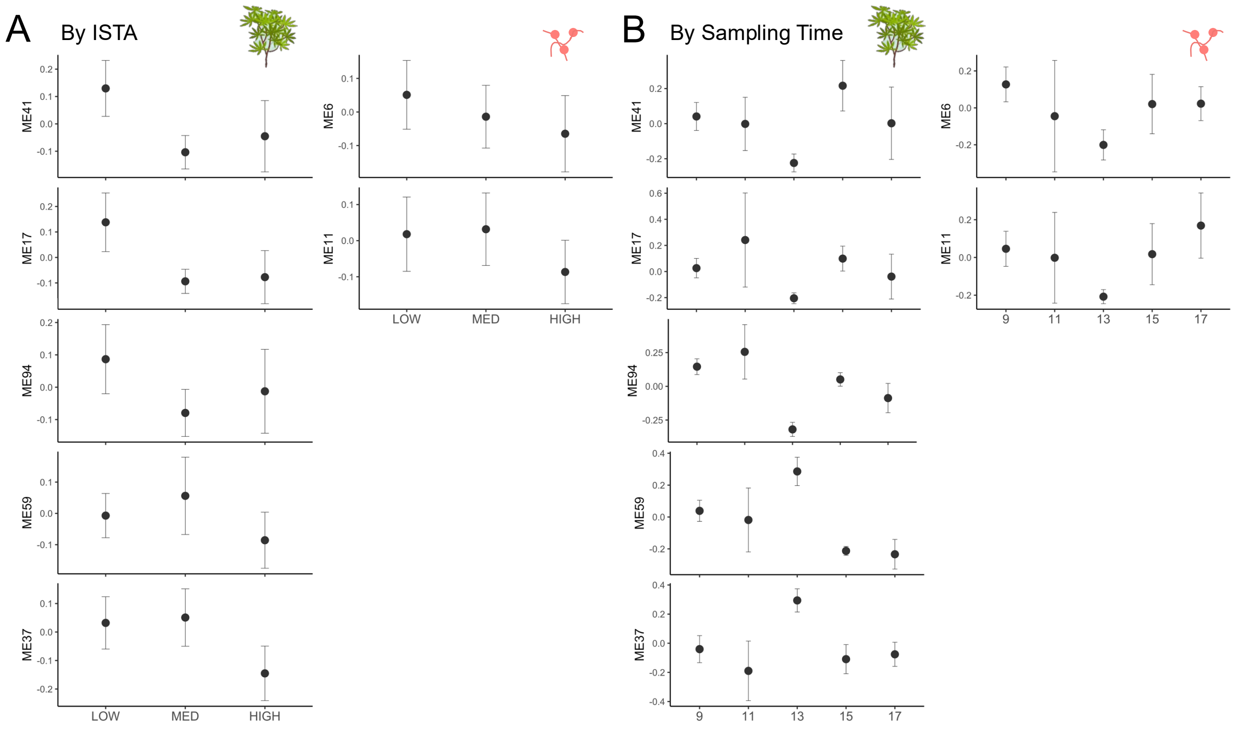

**Fig. S6** Soil properties data as (A) gradients in the field site (6 examples representing the lack of obvious explanatory gradients) and (B) statistical tests on correlation analyses on shoot and root biomass results.

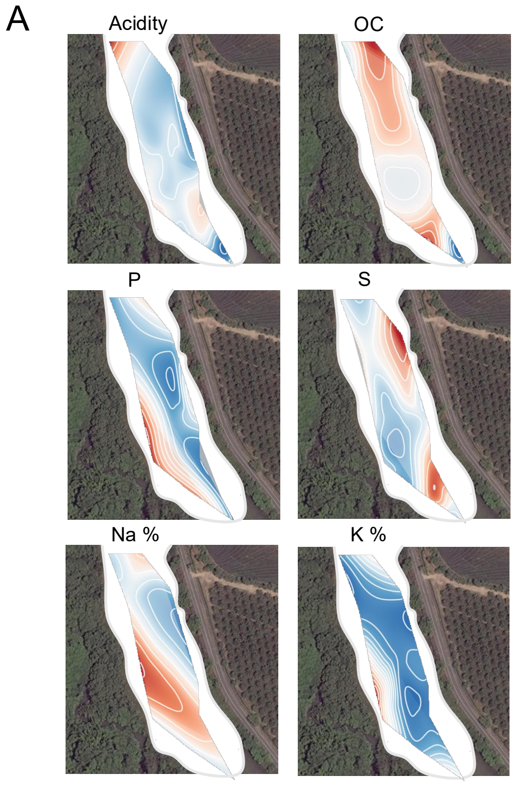


B

| Soil property | *p*-value correlation Shoot | *p*-value interaction Shoot:Isolate | *p*-value correlation Root | *p*-value interaction Root:Isolate |
|---|---|---|---|---|
| Sand | 0.06 | 0.54 | 0.52 | 0.33 |
| Silt | 0.16 | 0.20 | 0.45 | 0.44 |
| Clay | 0.13 | 0.53 | 0.64 | 0.25 |
| pH | 0.21 | 0.54 | 0.28 | 0.25 |
| Organic C (OC) | 0.06 | 0.26 | 0.12 | 0.81 |
| K | **0.01** * | 0.60 | 0.66 | 0.73 |
| Ca | 0.12 | 0.57 | 0.26 | 0.43 |
| Mg | 0.17 | 0.37 | 0.49 | 0.43 |
| Na | 0.05 | 0.42 | 0.20 | 0.75 |
| Acidity | 0.43 | 0.43 | 0.76 | 0.08 |
| Fe | 0.13 | 0.44 | 0.08 | 0.14 |
| Mn | **0.04** * | 0.96 | 0.59 | 0.32 |
| Cu | 0.08 | 0.64 | 0.22 | 0.13 |
| Zn | **0.03** * | 0.97 | 0.58 | 0.61 |
| B | 0.70 | 0.99 | 0.61 | 0.79 |
| P | **0.01** * | 0.68 | 0.10 | 0.64 |
| S | 0.77 | 0.44 | 0.96 | **0.03** * |
| Mg % | 0.79 | 0.72 | 0.31 | 0.22 |
| Na % | 0.19 | 0.41 | 0.11 | 0.44 |
| Al % | 0.35 | 0.45 | 0.56 | 0.12 |
| K % | **0.04** * | 0.77 | 0.57 | 0.52 |
| Ca % | 0.37 | 0.29 | 0.12 | 0.38 |

New Phytologist

**Fig. S7** Putative peaking times of the (A) plant Index of Symbiotic Transcriptional Activity (ISTA) and (B) *R. irregularis* presence in plant roots, as determined by an *Ri* housekeeping gene.

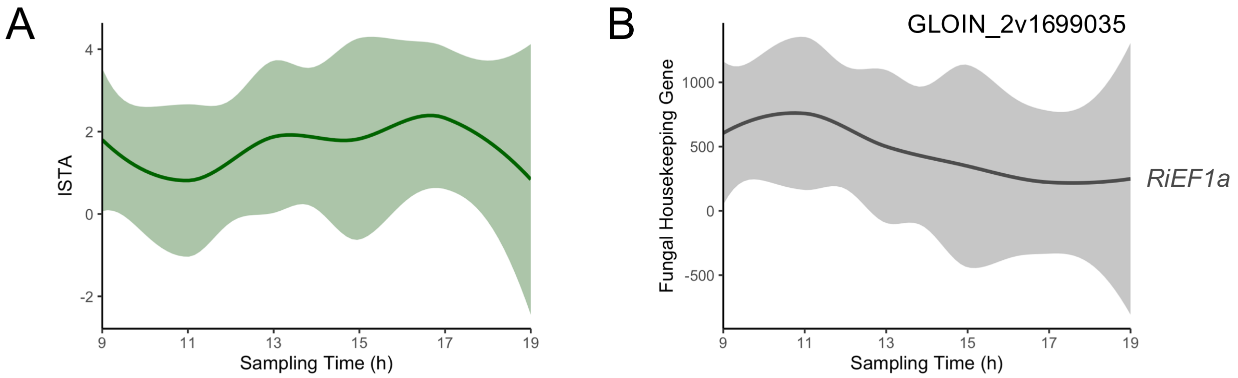

**Table S1** *R. irregularis* isolates used and studies where they were previously characterized.

| **Isolate type** | **Isolate ID** | **Previous characterization(s)** |
|---|---|---|
| Parent | A1 | Wyss *et al.* 2016; Savary *et al.* 2018 |
| Parent | A5 | Wyss *et al.* 2016; Savary *et al.* 2018 |
| Single-spore progeny of A5 | A5.2 | Robbins *et al.* 2021 |
| Single-spore progeny of A5 | A5.4 | Robbins *et al.* 2021 |
| Parent | B12 | Wyss *et al.* 2016; Savary *et al.* 2018 |
| Single-spore progeny of B12 | B12.1 | Robbins *et al.* 2021 |
| Single-spore progeny of B12 | B12.4 | Robbins *et al.* 2021 |
| Parent | C3 | Wyss *et al.* 2016; Savary *et al.* 2018 |
| Parent | C5 | Wyss *et al.* 2016; Savary *et al.* 2018 |
| Single-spore progeny of C2 | C2.5 | Robbins *et al.* 2021 |
| Single-spore progeny of C2 | C2.8 | Robbins *et al.* 2021 |

**Table S2** *M. esculenta* and *R. irregularis* clock gene IDs.

| Organism | Gene shorthand | Gene ID | Analysis notes |
|---|---|---|---|
| *M. esculenta* | *LHY* | Manes.01G239000.v7.1 | Averaged for relative |
| | *LHY* | Manes.05G015600.v7.1 | transcript abundance |
| | *PRR5* | Manes.01G239500.v7.1 | |
| | *PRR7* | Manes.04G027400.v7.1 | |
| | *TOC1* | Manes.06G089800.v7.1 | Averaged for relative |
| | *TOC1* | Manes.14G081500.v7.1 | transcript abundance |
| *R. irregularis* | *FRQ* | GLOIN_2v1575682 | |

New Phytologist

**Table S3** Metadata for the samples taken for transcriptomic analysis including retroactively determined sampling time, and Index of Symbiotic Transcriptional Activity (ISTA) as a continuous and categorical variable.

| RNAseq sample | Isolate Treatment | Block | Sampling Time | ISTA Continuous | ISTA Categorical |
|---|---|---|---|---|---|
| COL_FEB_01 | Z31 | 8 | 17:00 | 1.00 | MED |
| COL_FEB_02 | X35 | 8 | 15:00 | 1.97 | MED |
| COL_FEB_03 | Z31 | 12 | 9:00 | 0.02 | LOW |
| COL_FEB_04 | A1 | 4 | 11:00 | 0.02 | LOW |
| COL_FEB_05 | X39 | 3 | 17:00 | 0.03 | LOW |
| COL_FEB_06 | A5 | 3 | 19:00 | 0.98 | MED |
| COL_FEB_07 | X21 | 12 | 13:00 | 0.14 | MED |
| COL_FEB_08 | A1 | 8 | 11:00 | 0.03 | LOW |
| COL_FEB_09 | C3 | 10 | 13:00 | 0.83 | MED |
| COL_FEB_10 | B12ssp3 | 9 | 13:00 | 3.55 | HIGH |
| COL_FEB_11 | X39 | 2 | 13:00 | 0.70 | MED |
| COL_FEB_12 | A5ssp2 | 12 | 9:00 | 0.38 | MED |
| COL_FEB_13 | A5 | 12 | 11:00 | 0.14 | MED |
| COL_FEB_14 | C3 | 3 | 11:00 | 1.02 | MED |
| COL_FEB_15 | X21 | 8 | 15:00 | 1.67 | MED |
| COL_FEB_16 | A5ssp2 | 2 | 17:00 | 7.75 | HIGH |
| COL_FEB_17 | A5ssp2 | 9 | 11:00 | 3.68 | HIGH |

| | | | | | |
|---|---|---|---|---|---|
| COL_FEB_18 | B12ssp3 | 12 | 9:00 | 1.79 | MED |
| COL_FEB_19 | A1 | 5 | 17:00 | 3.10 | HIGH |
| COL_FEB_20 | X39 | 1 | 9:00 | 0.01 | LOW |
| COL_FEB_21 | C5 | 5 | 13:00 | 1.42 | MED |
| COL_FEB_22 | B12 | 7 | 13:00 | 0.01 | LOW |
| COL_FEB_23 | B12 | 12 | 17:00 | 2.24 | HIGH |
| COL_FEB_24 | C5 | 12 | 11:00 | 0.78 | MED |
| COL_FEB_25 | X21 | 5 | 13:00 | 6.47 | HIGH |
| COL_FEB_26 | C5 | 1 | 9:00 | 0.02 | LOW |
| COL_FEB_27 | B12ssp3 | 12 | 9:00 | 0.08 | MED |
| COL_FEB_28 | X35 | 5 | 15:00 | 3.63 | HIGH |
| COL_FEB_29 | B12 | 9 | 11:00 | 0.02 | LOW |
| COL_FEB_30 | C3 | 2 | 19:00 | 0.03 | LOW |
| COL_FEB_31 | Z31 | 10 | 9:00 | 3.37 | HIGH |
| COL_FEB_32 | A5 | 4 | 9:00 | 8.74 | HIGH |
| COL_FEB_33 | X35 | 2 | 15:00 | 0.02 | LOW |

**Table S4** *M. esculenta* gene IDs used to calculate the Index of Symbiotic Transcriptional Activity (ISTA) raw (continuous) values.

| Organism | Gene shorthand | Gene ID |
|---|---|---|
| *M. esculenta* | *LEC5.7* | Manes.12G124400.v7.1 |
| | *RAM2* | Manes.01G193000.v7.1 |
| | *GLP2* | Manes.04G002400.v7.1 |
| | *GST1* | Manes.01G123300.v7.1 |
| | *HA1* | Manes.05G053600.v7.1 |

**Table S5** Information on the ranking group divisions for the CEMiTool analysis.

| Ranking group | Root biomass range | Mean ± SE | Replicates |
|---|---|---|---|
| 1 | 5.655 – 8.888 kg | 6.774 ± 0.402 | 7 |
| 2 | 4.696 – 5.255 kg | 4.926 ± 0.083 | 8 |
| 3 | 3.165 – 4.515 kg | 4.069 ± 0.167 | 8 |
| 4 | 0.995 – 2.990 kg | 2.061 ± 0.297 | 7 |

**Table S6** Details on genes in the modules of interest from the CEMiTool analysis (NC, M8, M4).

*Submitted as an Excel file.*

**Table S7** Details on genes in the final 7 WGCNA modules of interest.

*Submitted as an Excel file.*
